# Distinguishing Felsenstein zone from Farris zone using neural networks

**DOI:** 10.1101/822288

**Authors:** Tamara Drucks, Alina F. Leuchtenberger, Sebastian Burgstaller-Muehlbacher, Stephen M. Crotty, Heiko A. Schmidt, Arndt von Haeseler

## Abstract

Maximum likelihood and maximum parsimony are two key methods for phylogenetic tree reconstruction. Under certain conditions, each of these two methods can perform more or less efficiently than the other. We show that a neural network can efficiently distinguish between four-taxon alignments that were evolved under conditions conducive to long-branch attraction, or long-branch repulsion. The feedback from the neural network can be used to select the most efficient tree reconstruction method yielding increased accuracy, when compared to a rigid choice of reconstruction methods. When applied to the contentious case of Strepsiptera evolution, our method agrees with the current scientific view.

## Introduction

The phylogenetic artefact known as long-branch attraction (LBA) was first brought to attention in Felsenstein’s seminal paper (Felsenstein, 1978), and subsequently elaborated on by Hendy and Penny (1989). They describe LBA succinctly, as the phenomena whereby “two long-branched, non-sister taxa are grouped together, rather than with their shorter-branched sister taxa” when performing inference by maximum parsimony (MP). In recent years, LBA has, to some extent, lost its precise meaning (Bergsten 2005; Sanderson et al. 2000), but for the sake of clarity we will use the term LBA *sensu stricto*. The simplest form of LBA occurs in a four-taxon tree, with two long branches and three short branches as displayed in Figure 1a-b. When the long branches are substantially larger than the short branches, MP is statistically inconsistent. That is, it will fail to reconstruct the true evolutionary history, instead grouping the taxa with the long branches in one clade, even with infinite sequence length. On the other hand, in the same scenario, maximum likelihood inference (ML) is statistically consistent. Given infinite sequence length, ML will recover the true tree, assuming the correct model of sequence evolution. Huelsenbeck and Hillis (1993) coined the term “Felsenstein zone” to refer to parameter combinations for which MP was inconsistent.

**Fig. 1:**
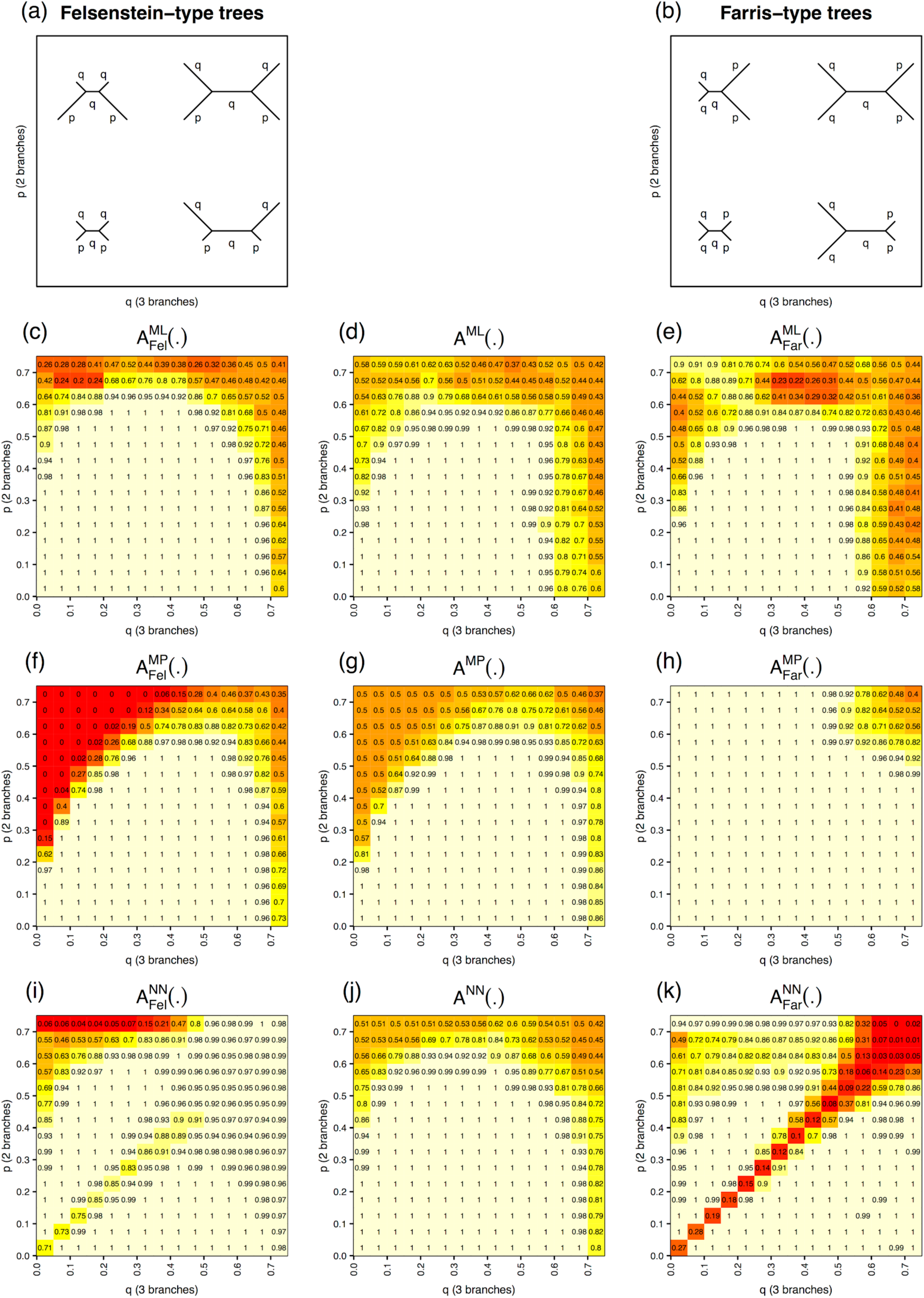
Visualization of Felsenstein-type (a) and Farris-type tree topologies (b); accuracies of phylogenetic reconstruction using ML (c-e), and MP (f-h); and to infer the tree-type with the NN (i,k). The accuracy to reconstruct the tree using the integrated strategy using NN, MP and ML (j).

However, the asymptotic behaviour of the reconstruction methods is of limited relevance, given the finite sequence lengths of biological data. In practice, the principal concerns relate to whether LBA is a relevant issue with biological datasets; and if so, the efficiency of tree reconstruction methods (Hillis et al. 1994) in the presence of LBA.

While LBA was conclusively demonstrated using simulations by the late 1980s, considerable debate centred on whether this artefact was only of theoretical interest or could actually manifest in biological datasets. The first biological example of LBA was introduced by Carmean and Crespi (1995), arguing that the LBA artefact was responsible for the placement of Strepsiptera as sister to Diptera among insects. Their conclusion was challenged by Huelsenbeck (1997), who discussed the lack of evidence for LBA and suggested a simulation-based method to detect whether branches are long enough to be attracted by parsimony. He also pointed out the need for an unbiased reconstruction method that was not affected by LBA. Subsequently, Siddall (1998) asserted that LBA was not a significant issue in practice, showing that a similar effect in the opposite direction could also be demonstrated through simulation on four taxa. By constructing the true tree such that the long branches were in fact sisters, Siddall showed that parsimony was more efficient than likelihood in reconstructing the phylogeny (although likelihood methods remained consistent). He coined the parameter combinations for which parsimony was more efficient than likelihood, the “Farris zone”. Further, he reasoned that since the truth was unknowable in real datasets, when faced with discordance between parsimony and likelihood methods, it would be impossible to tell whether two long branches grouped together had been resolved correctly or not. This argument has held firm ever since.

Previous attempts to use neural networks to classify Farris-/Felsenstein-type trees have shown mixed results (Suvorov et al. 2019; Zou et al. 2019). We demonstrate here that a simple, feedforward neural network can distinguish between alignments derived from a Felsenstein-type tree (two long branches in a four-taxon tree separated by a short internal edge) and a Farris-type tree (two long branches forming a cherry). Feedback from the network can then be used to inform reconstruction method selection.

## Methods

### Neural network

Neural networks are computing systems that attempt to emulate particular features of the biological brain of sentient beings, namely the ability to learn from experience. A neural network is trained by inputting large amounts of data with known output values. The network is then exposed to new data, which it classifies based on its training. Inspired by recent advances in the application of neural networks to a wide range of problems, in particular its strength in pattern recognition (Goodfellow 2016 and references therein), we employ a feedforward neural network (e.g. Nielsen 2015) to classify multiple sequence alignments according to their tree types (i.e. whether they are Felsenstein-type or Farris-type).

For a detailed overview of our network architecture, see Supplementary Table S1.

### Data preprocessing

To encode multiple sequence alignments into a suitable format for the neural network, we computed the site-pattern frequencies of each alignment. For four taxa, 256 unique site-patterns exist for the four-letter DNA alphabet. When using the JC-model (Jukes and Cantor 1969), the 256 patterns collapse to 15 distinct pattern-categories due to the symmetries in the substitution model. These are xxxx, xxxy, xxyx, xyxx, xxyy, xxyz, xyxy, xyyx, xyyy, xyxz, xyyz, xyzx, xyzy, xyzz, xyzw, where x, y, z and w denote different nucleotides. For each parameter combination (p,q) the probabilities of the 15 patterns can be computed analytically (Felsenstein 2004, pp. 111), thus generating a multinomial distribution MD(p,q).

When carrying out the simulation-based training of the neural network, we are of course aware which taxon belongs to which branch length (i.e., which are from short branch lengths and which are from long). When analysing empirical data we do not have this luxury, and so the network must be able to accurately classify alignments independent of the order of the taxa. To achieve this, we permuted the training data such that each simulated alignment was presented to the network 24 times, once for each different ordering of the four taxa.

### Data simulation

To keep everything simple and well defined, we assumed the JC-model of sequence evolution for our initial experimental setup. Figure 1a-b shows the two trees that have a parameter p for two branches and a parameter q for three branches, for different (p,q) combinations. The parameters p and q describe the probabilities of observing a substitution at a particular site along that branch.

We will distinguish between Felsenstein/Farris zone and Felsenstein-/Farris-type trees. The Felsenstein zone is defined by the classical definition following Felsenstein (1978). However, the Farris zone is not unambiguously defined for ML-inference as its boundary depends on the length of the sequence alignment. To avoid this ambiguity, we simulate four-taxon trees for the full range of parameter (p,q) combinations. It should therefore be noted that the trees are indistinguishable if p=q and will switch from two long branches to three long branches as p<q.

We independently varied p and q from a minimum of 0.005 to a maximum of 0.745 at increments of 0.01. This created a total of 75×75 = 5625 different parameter combinations. For each of these parameter combinations, the probability of observing each of the 15 pattern-categories was computed, forming a multinomial distribution, MD(p, q). From this multinomial distribution, 1000 pattern-categories were drawn randomly to simulate a sequence alignment of length 1000 nucleotides. For each combination of p and q 1000 training alignments were simulated.

Test alignments were generated for a more sparsely populated grid of *p* and q combinations. The parameters were varied between 0.025 and 0.725 at increments of 0.05, creating a total of 15×15=225 parameter combinations. For each combination of *p* and *q* we used the program Seq-Gen (Rambaut and Grassly 1997) to simulate 200 different multiple sequence alignments of length 1000 nucleotides. These alignments were processed to convert them into vectors containing the relative pattern-category frequencies. The pattern-category frequencies served as input for the neural network and the classical unweighted MP analysis, while the alignments served as input for phylogenetic inference using ML with IQ-TREE (Nguyen et al. 2015).

### Analysis of real data

For our experimental test data we revisit the well-known problem of placing Strepsiptera in the phylogenetic tree (Carmean and Crespi 1995; Whiting 1997; Huelsenbeck 1997) of insects. The dataset consists of 18S ribosomal DNA sequence data for 13 insect species. As described by Huelsenbeck (1997), we removed all sites with gaps and unknown DNA characters. We subsampled sets of four sequences (quartets) from the original alignment, strategically selecting taxa such that each quartet would be relevant to the placement of Strepsiptera. We constructed 24 such quartets. Each quartet comprised one sequence each of the four taxonomic groups: Strepsiptera (1), flies (2), beetles (2) and Holometabola (6). For each quartet, we estimated branch lengths for Felsenstein-like (Strepsiptera grouped with beetles) and Farris-like (Strepsiptera grouped with flies) topologies. We repeated this parameter estimation procedure for twelve different substitution models: JC (Jukes and Cantor 1969), K2P (Kimura 1980), F81 (Felsenstein 1981), HKY (Hasegawa et al. 1985), TN (Tamura and Nei 1993), GTR (Tavaré 1986), and their respective Gamma-variants (+G; Yang 1994). Model parameters were estimated using the full alignment with and without outgroup sequences. ML branch lengths and model parameters were inferred using IQ-TREE.

The process described above yielded 1152 different quartet tree parameterisations (24 quartets x 2 topologies x 12 models x with/without outgroup). For each of these quartet parameterisations we simulated 10000 alignments using Seq-Gen to be used as training data. We simulated an additional 10 alignments per parameterisation for network validation purposes. The extension of our simulation study to empirical data required all 256 site-pattern frequencies to be considered. This necessitated changes to the network architecture, which are detailed in Supplementary Fig. S1. As test data we extracted the relevant 24 quartet sub-alignments from the biological alignment described above. These were summarised into site-pattern frequency vectors which then served as input for the neural network.

## Results & Discussion

### Simulation Study

Across most of the parameter space, the neural network was able to distinguish whether an alignment originated from Felsenstein-like or Farris-like trees (Fig. 1i,k). We now can use the network as an upstream instance to suggest the preferred reconstruction method. The neural network, thus, acts as an impartial judge, deciding whether using MP or ML is more adequate to reconstruct the phylogenetic tree.

There are two main regions of concern for the network. Naturally, the network cannot distinguish the data if p=q as the trees are identical (cf. the diagonal in Fig. 1k) and so it’s performance is poor in this area. Additionally, the network performs poorly for Felsenstein-type trees with very high p and very low q (see Fig. 1i). However, these two areas of concern for the network are of little practical consequence, as MP and ML perform similarly in these regions (Figs. 1c,e,f,h).

More formally the accuracies shown in Figure 1c-k can be computed as follows:

With 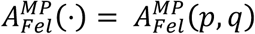 we denote the accuracy of MP under a Felsenstein-type tree with branch lengths *p* and *q*. The other definitions are analogous.

Figure 1d and 1g show the total accuracies for ML and MP independent of the mode of evolution which can be computed as

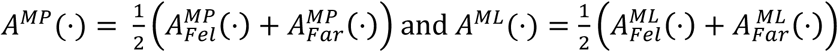

(See Supplementary Material).

Finally, we use the information provided by neural network for an integrated strategy that selects the more reliable tree reconstruction algorithm. The accuracies of this strategy can be calculated by

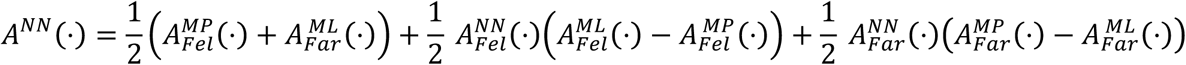

where the derivation of the formula can be found in Supplementary Material.

Figure 1j shows the results of the boosted performance using this equation. Finally, it provides further insights: The first summand, the average accuracy of MP and ML under the unfavourable evolutionary scenario, i.e. (p,q) are from the Felsenstein/Farris zone, can only be improved if the sequence length *l* increases. For *l* → ∞ we obtain

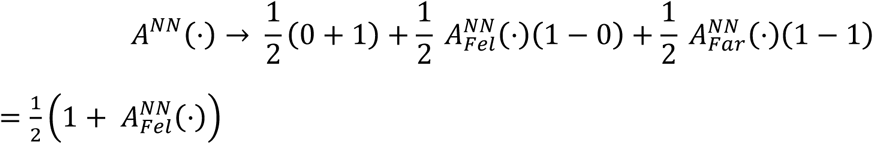

if (p,q) are from the original Felsenstein/Farris zone. We can improve the overall accuracy by training the network to reliably detect the Felsenstein-type trees, thus 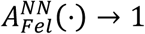. If (p,q) are from the non-zone and *l* → ∞ then MP and ML will identify the true tree and the decision of the neural network does not matter. Accordingly, the neural network shows better accuracies for Felsenstein-type trees than for Farris-type trees.

To assess the impact of the information available we tested the neural network with alignments of various lengths up to 10000bp. With increasing sequence length, the accuracy of the network improves.

### Analysis of real data

The network trained for use with the empirical data was also able to successfully distinguish between Felsenstein-like and Farris-like trees. On the simulated validation data the network correctly classified 96.08% of the Felsenstein-type trees and 96.52% of the Farris-type trees. When we tested the 24 quartet alignments taken from the original data of Carmean and Crespi, the results were unanimous. For all 24 quartets the network predicted that these data had evolved on a Felsenstein-like tree. Therefore, it suggests a placement of Strepsiptera distant from the flies, which is in accordance to Niehuis et al. (2012) and Boussau et al. (2014).

## Conclusion

We have explored a well-known Achilles Heel of phylogenetic inference, LBA, and demonstrated that neural networks can be employed to inform the choice of phylogenetic inference method, potentially improving accuracy and increasing efficiency. Further, we show that the application of our method to a dataset that has been contentious in the literature, with some considering it an example of LBA and others not, yields results consistent with the currently accepted phylogeny. This initial study illustrates the potential of neural networks to be applied to the tree inference problem. Our approach suggests that in the face of topological discordance among competing inference methods, machine learning techniques may be able to point towards the underlying biological truth. Moreover, we are able to reconstruct simple trees using neural networks.

In this study we have only scratched the surface of the potential of deep learning approaches to be utilized in the field of phylogenetic inference. We think it is timely to share the results with the scientific community, to promote the application of deep learning approaches to an array of interesting phylogenetic questions.

## Acknowledgements

This work was supported by the Austrian Science Fund (FWF DOC 32-B28 to A.L. and A.v.H.).

A.v.H. also thanks the Medical University of Vienna and the University of Vienna for their support. We thank the *Internet Archive* (archive.org) for providing the possibility to recover the lost original dataset of Carmean and Crespi (1995).

The authors declare they have no competing interests.

## Supplementary Material

### Network Architecture

The trained networks as well as the used training and test data can be found at GitHub (https://github.com/Cibiv/zone-net). An overview of the architecture and hyperparameters of the networks is presented in Tab. S1.

**Table S1:** Architecture and hyperparameters of the neural network for simulated alignments using the Jukes-Cantor model and the neural network data used on Strepsiptera data of Carmean and Crespi (1995)

| Network for | simulated Jukes-Cantor data | Strepsiptera data |
| --- | --- | --- |
| Number of nodes in layers | 15, 64, 128, 256, 512, 1024, 408, 208, 96, 1 | 256, 382, 512, 892, 1024, 408, 324, 208, 96, 1 |
| Transfer function (hidden layers) | ReLU | ReLU |
| Activation function (output layer) | Sigmoid | Sigmoid |
| Weight initialization | Xavier initialization (Glorot 2010) | Xavier initialization (Glorot 2010) |
| Bias initialization | Zero initialization | Zero initialization |
| Learning rate | 0.0001 | 0.0001 |
| Batch size | 32 | 32 |
| Cost function | Sigmoid cross-entropy | Sigmoid cross-entropy |
| Optimizer | Adam (Kingma and Ba 2015) | Adam (Kingma and Ba 2015) |
| Epochs trained | 2 | 4 |

### Total accuracies of maximum likelihood, maximum parsimony and the integrated strategy

We denote the accuracy of maximum parsimony (MP) under Felsenstein-type trees with branch lengths p and q by 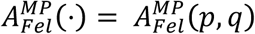. Analogously, we define the accuracy of MP under Farris-type trees as well as the accuracy of maximum likelihood (ML) and the neural network (NN).

We denote with *x* the probability that the alignment follows a Felsenstein-type tree. Then we can compute the total accuracy of MP independent of the mode of evolution as

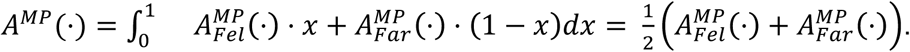

Similarly, we obtain the total accuracy of ML as

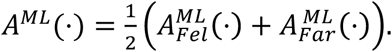

The total accuracy of the integrated strategy is then computed by integrating the network’s accuracy and the accuracy of the proposed methods over the probability x that the alignment originated of a Felsenstein-type tree:

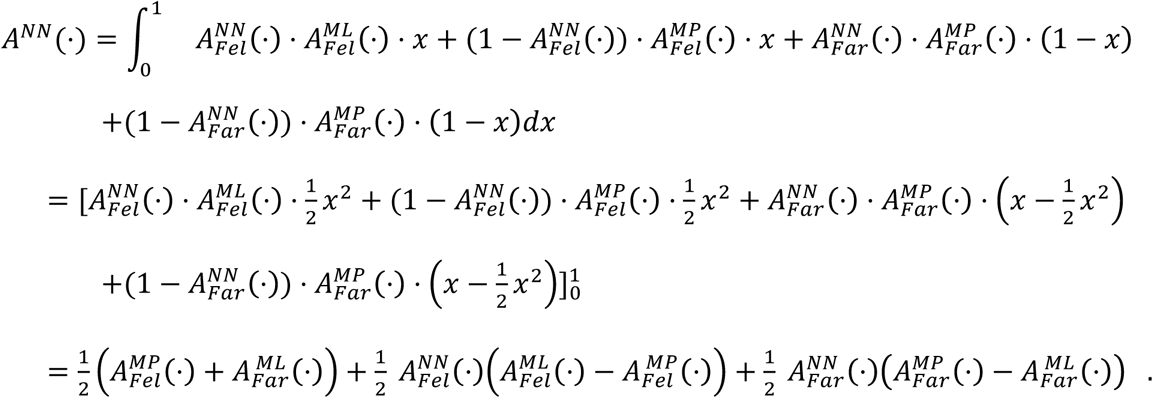

